# Applications of AlphaFold beyond Protein Structure Prediction

**DOI:** 10.1101/2021.11.03.467194

**Authors:** Yuan Zhang, Peizhao Li, Feng Pan, Hongfu Liu, Pengyu Hong, Xiuwen Liu, Jinfeng Zhang

## Abstract

Predicting structures accurately for natural protein sequences by DeepMind’s AlphaFold is certainly one of the greatest breakthroughs in biology in the twenty-first century. For designed or engineered sequences, which can be unstable, predicting the stabilities together with their structures is essential since unstable structures will not function properly. We found that experimentally measured stability changes of point mutations correlate poorly with the confidence scores produced by AlphaFold. However, the stability changes can be accurately predicted using features extracted from the representations learned by AlphaFold, indicating greater generalizability of AlphaFold to designed or engineered sequences than previously thought. We then used AlphaFold to validate our previously developed protein design method, ProDCoNN, that designs sequences to fold to target protein structures given only the backbone structure information of the target proteins. We showed that ProDCoNN was able to design sequences that fold to structures very close to target structures. By combining a modified ProDCoNN, AlphaFold, and sequential Monte Carlo, we designed a novel framework to estimate the designability of protein structures. The designability of a protein structure is defined as the number of sequences, which encode the protein structure, and is an indicator of the functional robustness of proteins. For the first time, we estimated the designability of a real protein structure, chain A of FLT3 ligand (PDB ID: 1ETE) with 134 residues, as 3.12±2.14E85.

## Introduction

Protein structure prediction (PSP) has been one of the most challenging problems in computational biology^1^. It is also a problem whose solution will have a profound impact in many areas of biology and biomedical sciences. Not surprisingly, the problem has attracted researchers from many different disciplines for half a century since it was originally proposed^2^. Critical Assessment of Structure Prediction (CASP) was initiated in 1994 to provide a blind test of methods for PSP^3,4^, which has played a key role for advancing PSP methods. Except the early years of CASP with some substantial progress, the field had come to a standstill for quite some years, until in 2018 when DeepMind joined the game with their deep learning powered method, AlphaFold, which substantially improved the prediction accuracy in CASP13^5^. And in CASP14 held in 2020, AlphaFold2^6^ (we will call it AlphaFold in the rest of the paper) has pushed the numbers so much so that the organizers of CASP declared that the PSP problem has been finally solved^4^. Since then, DeepMind team has applied AlphaFold to predict more than 350,000 protein structures in human and other species, and released the structures for biological community to use freely^7^. These predicted protein structures will help biologists infer the functions of these proteins to better understand the mechanisms of the biological processes or diseases they are involved in. In a follow-up study^8^, and also by Baker and co-workers who developed RoseTTAFold using a deep learning framework inspired by AlphaFold^9^, it has been shown that the protein-protein interaction problem can be considered as a protein structure prediction problem by putting two or more protein chains together to predict their complex structures.

One important and related question unanswered by AlphaFold is whether the predicted structures are stable enough for their functions. For natural sequences, it may not be a serious issue since most of them fold to stable structures. However, for a designed or engineered sequence, which can be unstable, AlphaFold predicts a structure regardless whether it is stable or not. This is because AlphaFold was trained using only sequences that can fold to stable three-dimensional structures. Predicting a structure whose stability is poor can be misleading. Strictly speaking, predicted stability should come with predicted structure to be a meaningful prediction. As of now, AlphaFold was considered as not suitable for modeling point mutations^10^. This is consistent with our finding that the confidence scores of AlphaFold correlates poorly with the experimentally measured stability changes for point mutations (Fig 1). There are two possible explanations for this observation. Firstly, AlphaFold may have learned some sophisticated patterns in the natural protein sequences and structures to predict structures accurately while bypassing the learning of any energetics of proteins such as the complex physical interactions among the atoms. If this is the case, then it may not work well for unseen cases such as point mutations due to the bias in the training data. In the training data, similar sequences always have very similar and stable structures since unstable ones will not be in PDB, and will not be part of the training data. The second explanation is that AlphaFold actually learns some energetics of proteins using the sophisticated neural network architecture, which made it possible to predict structures accurately. Since it never used any stability data, the model was not trained to produce any outputs that correlate with stabilities. The confidence scores were produced for a totally different purpose. If this is the case, then it may be possible to extract some representations from the intermediate outputs of AlphaFold, and take them as input with the stability data of point mutations as output to train a model to predict the stability changes of point mutations. This was one of the key hypotheses we tested in this study.

**Fig 1.**
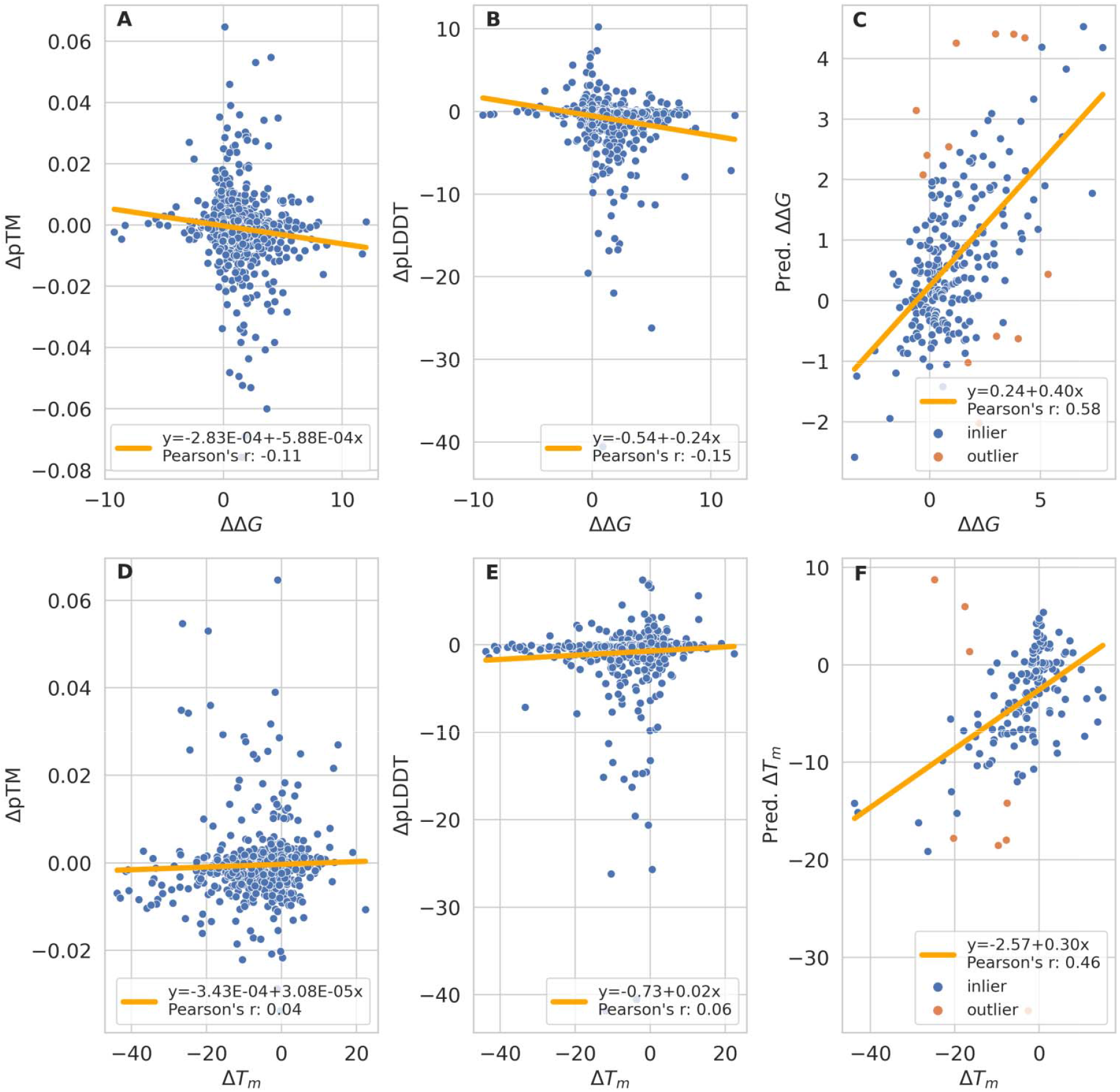
Modeling the point mutations using AlphaFold. **A**. ΔpTM for whole sequences between wild type and mutant vs. ΔΔ*G* of the mutation; **B**. ΔpLDDT for the mutated residue between wild type and mutant vs. ΔΔ*G* of the mutation; **C**. Predicted ΔΔ*G* using representations learned by AlphaFold vs. experimentally measured ΔΔ*G*; **D**. ΔpTM for whole sequences between wild type and mutant vs. ΔΔ*T*_*m*_ of the mutation; **E**. ΔpLDDT for the mutated residue between wild type and mutant vs. ΔΔ*T*_*m*_ of the mutation; **F**. Predicted ΔΔ*T*_*m*_ using representations learned by AlphaFold vs. experimentally measured ΔΔ*T*_*m*_.

Indeed, our study showed that the representations learned by AlphaFold during the structure prediction process can be used to predict stability changes of point mutations quite accurately. We obtained the state-of-the-art performance using a relatively simple model with a smaller training dataset compared to existing methods. This interesting finding indicates that it is likely that AlphaFold has learned the fundamental physical principles of proteins in some forms, which make it generalizable to unseen situations – sequences not well-represented in the training data. With this observation, in this study, we explored several applications of AlphaFold beyond predicting structures for natural sequences.

We then used AlphaFold to predict the structures for the sequences designed to fold to target structures using a protein design method we developed recently, ProDCoNN^11^. ProDCoNN used a deep neural network architecture to model the local three-dimensional environment of individual residues. It achieved the best performance for the inverse protein folding (IPF) problem tested on benchmark datasets. We found that some designed sequences could fold to structures quite close (<4Å RMSD) to the target structures while others cannot (as predicted by AlphaFold). AlphaFold can thus be used to select promising sequences designed for an IPF problem. We then propose a new framework by combining AlphaFold, ProDCoNN, and sequential Monte Carlo (SMC) to estimate the designability of a given protein structure. The designability of a given protein structure is defined as the number of sequences that encode the structure^12,13^. A sequence encodes a structure if the structure is very close to the native structure of the sequence. Designability has been proposed to contribute to protein structure and function robustness^12^, and it has also been shown to correlate with the relative frequencies of disease-causing proteins in a fold^13^. In this study, we used RMSD to measure the similarity between two structures and selected a reasonable cutoff value to define designability. For the first time, we have estimated the designability of a real protein structure, chain A of FLT3 ligand (PDB ID: 1ETE) with 134 residues, as 3.12±2.14E85.

It is well-known that natural proteins can tolerate many mutations and at the same time certain point mutations can severely destabilize a protein. Clearly, for a sequence to fold to a given structure, some positions can only accommodate certain types of amino acids. If we call such constraints as the minimum folding elements (MFEs) of a protein structure, a fundamental question is: given a protein structure what are its MFEs? A simple model to describe the MFEs of a structure would be position specific weight matrix or position weight matrix^14,15^. A reasonable way to estimate it would be to first find all the homologous sequences of the protein sequence for which we would like to find MFEs. We can then perform a multiple sequence alignment (MSA) for all these sequences to identify conserved residues for each position of the sequence. However, this may give a very biased estimation of the true MFEs of the protein structure, because some conserved positions may be conserved for functions, not for structure stability. In addition, the sequences we used in the MSA may not be an unbiased sample from all the sequences that can fold to a target structure. In this study, we show that the conservation pattern obtained using an MSA for homologous protein sequences is indeed very different from the pattern obtained from a set of foldable sequences predicted by AlphaFold. To identify the MFEs of a given structure, one can perform computational mutagenesis experiments using AlphaFold to sample the sequences that can fold to the structure and then perform an MSA. Such studies may also shed light on the fundamental principles of protein folding.

## Results

### Predicting stability changes of point mutations

We first applied AlphaFold to the point mutation dataset to see whether there is any relationship between the confidence scores it outputs and the experimentally measured stability changes. For a protein, *p*, we used AlphaFold to predict structures for both its wild type sequence and single-point mutant to generate two structures *S*_*p,w*_ and *S*_*p,m*_, respectively.

We examined the correlation between the stability changes and confidence scores output by AlphaFold. There are two different measures of stability changes for the mutants from FireProtDB database, ΔΔ*G* and Δ*T*_*m*_. There are also two different confidence scores (CSs) for the predicted structures, predicted TM-score, pTM, and predicted LDDT value for the mutated residue, pLDDT. It is more meaningful to look at the changes in pTM and pLDDT between *S*_*p,w*_ and *S*_*p,m*_. We plotted the scatter plots for all four possible pairs between stability changes and CS changes upon mutation (Fig 1A, 1B, 1D, 1E). None of the pairs showed any significant correlations.

We then explored the possibility of using the representations learned by AlphaFold to predict the stability changes of point mutations. We implemented a simple multilayer perceptron regression model for ΔΔ*G* (or ΔΔ*T*_*m*_) prediction (See Methods for details). We first used AlphaFold to predict structures of both wild type and mutant sequences. We then extracted the feature vectors from the position of the mutated residue from the “single representation” of the AlphaFold models for both wild type and mutant sequences as model input. We used 10-fold cross validation and the Pearson’s correlation coefficient between predicted ΔΔ*G* and experimental ΔΔ*G* is 0.58, which is significantly higher than those observed between the confidence scores and experimental ΔΔ*G*. It is also slightly higher than the state-of-the-art performance achieved by a recent deep learning method^16^. Fig 1C (ΔΔ*G*) and 1F (ΔΔ*T*_*m*_) show the scatter plot between predicted stability changes and experimental measured stability changes from 10-fold cross-validation.

### Predicting structures for sequences designed for inverse protein folding (IPF)

We next used AlphaFold to predict structures for sequences designed to fold to certain target protein structures. These sequences are significantly different from any of the natural sequences. Designing sequences that fold to a given protein structure is also called the inverse protein folding (IPF) problem. We selected several proteins from different fold classes from SCOPe database^17^. The sequences were designed using a modified ProDCoNN^11^ (see Method and Data for details). The similarities between designed sequences and the corresponding wild type sequences range between 16-36%. Fig 2 shows the predictions from AlphaFold for 4 different proteins. We can see that a significant number of the designed sequences are predicted to fold to relatively small RMSDs compared to the target structures. Our result indicates that AlphaFold can be used to select promising foldable sequences and filter out sequences that are not foldable (See Methods for the definition of foldable sequences).

**Fig 2.**
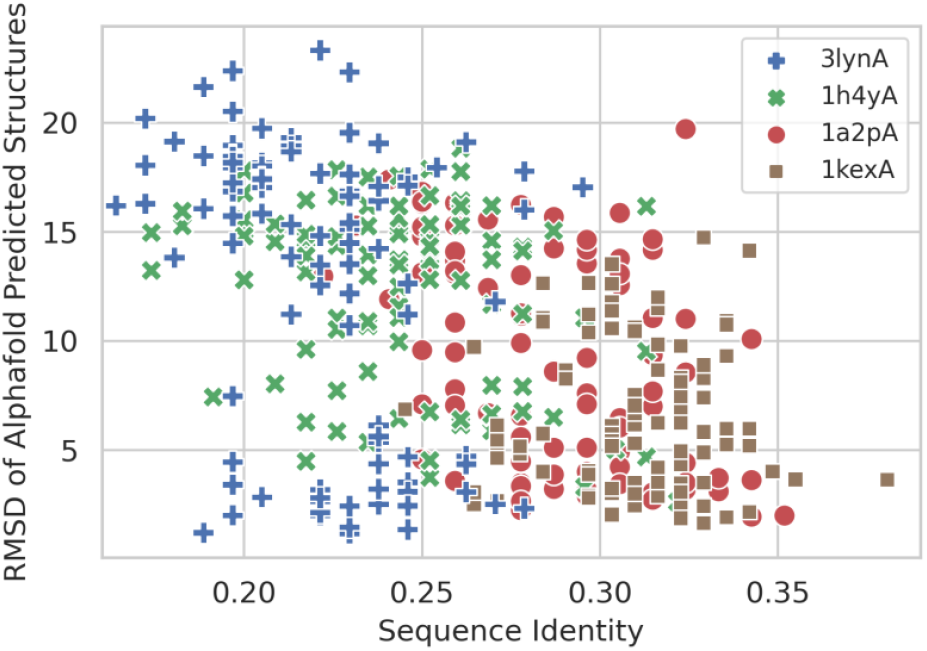
The RMSD of AlphaFold predicted structures to the target structures vs. the sequence identities of the designed sequences for 4 proteins. Significant numbers of sequences designed by ProDCoNN were predicted to fold to structures with relatively small RMSD to the target structures. The sequence identities were calculated by comparing to the sequences of the target structures. There is a significant negative correlation between RMSD and sequence identity across different proteins. But for individual proteins, the correlation is very weak. Clearly, the sequence identity is not the key factor for being foldable.

### Estimating the designability of real proteins

To estimate the designability of a protein structure, we first used the sequential Monte Carlo (SMC) strategy (See Methods for details) to sample a number of sequences for the protein structure. sampled sequence using SMC has a weight, which is updated recursively as follows: where *w*_*t*_ is the weight at step *t, w*_*t-*1_ is the weight at step *t*-1 and *p*_*t*_ is the probability the actual amino acid type at step *t* was sampled. The designability can then be estimated using the equation: –, where *D* is designability, *w*_*i*_ is the weight of sequence *i*, and *n* is the total number of foldable sequences.

In Fig 3, we estimated the designability for five proteins with lengths range from 89 to 166. We found that there is a general positive correlation between the length of a protein and its designability. However, length is not the only factor determining the designability. For example, protein 1ete chain A (1eteA) has 134 residues, and protein 2a2l chain A (2a2lA) has 145 residues, but 1eteA has significantly higher designability than 2a2lA. As far as we know, this is the first time designability was estimated for real proteins under very reasonable assumptions. For example, the designability of chain A of FLT3 ligand (PDB ID: 1ETE) with 134 residues was estimated as 3.12±2.14E85.

**Fig 3.**
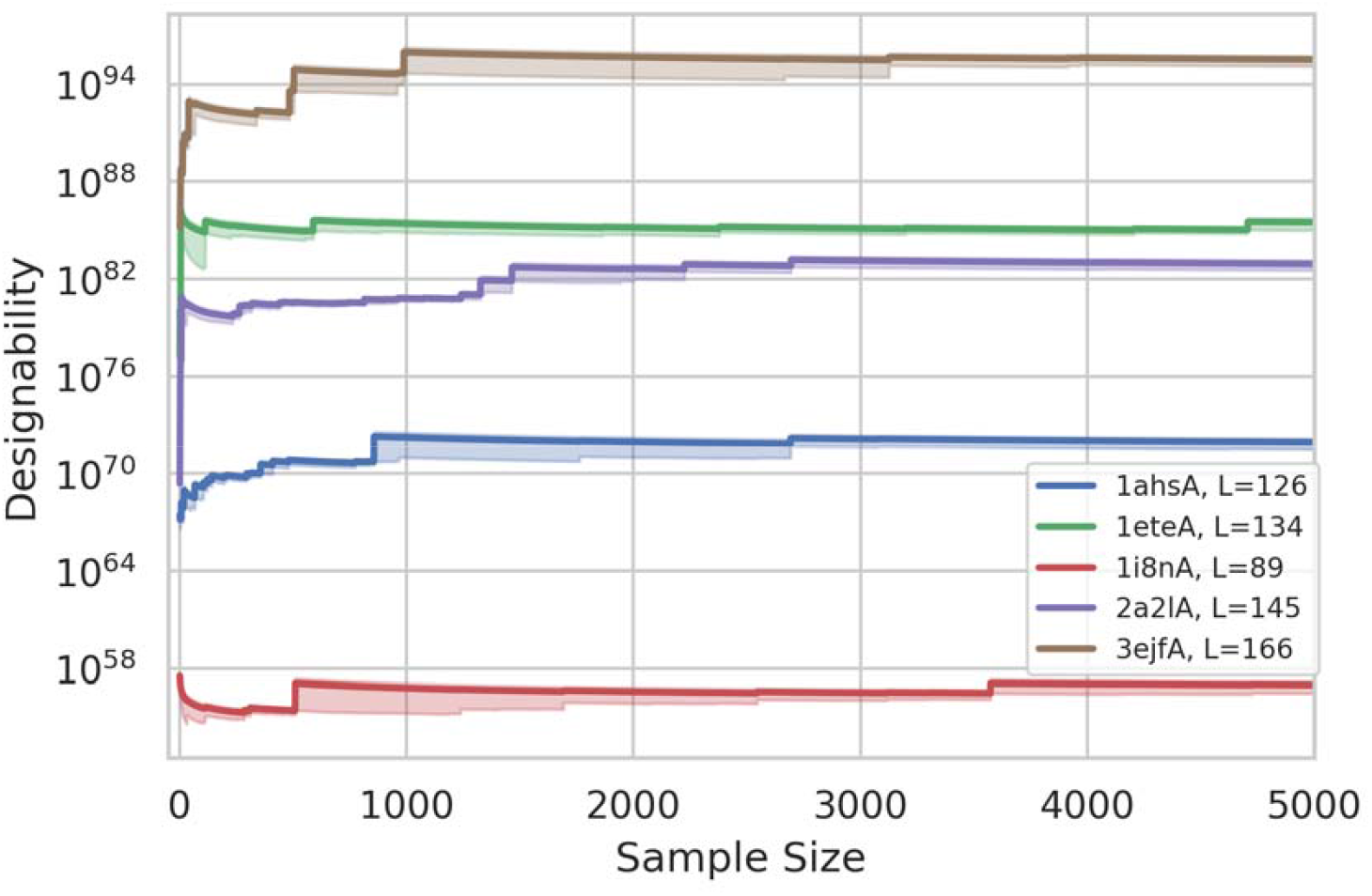
Designability for five proteins. The lengths, *L*, of the proteins are shown in the legend. We sampled 5000 sequences for each protein and estimated the designability for each sample size from 1 to 5000 to get the running estimated designability. The error bands are also shown for each estimation.

### Characterizing foldable sequences

For designed sequences, some are predicted to be foldable, while others are not. Studying the foldable sequences may reveal the key residues important for the folding and stability of a protein structure. Fig 4 shows two sequence logos, one plotted using foldable and one using homologous sequences obtained from multiple sequence alignment (MSA) for protein 1a2p. Clearly, despite significant similarity in many positions, the foldable sequences showed marked differences from homologous sequences at certain positions. This indicates that those conserved residues among the homologous sequences may be important for functions instead of stability. The comparison may shed light on potential point mutations that may increase the stability of the protein. It is worth noting that the MSA of 1a2p has much stronger conservations than the foldable sequences. The average entropy for MSA and foldable designed sequences are 0.813 and 1.367, respectively. The numbers of residues with entropy smaller than 1 for MSA and foldable sequences are 65 and 29, respectively. That indicates that the natural sequences may have only explored part of the foldable sequence space for this protein structure. The foldable sequences are more conserved than MSA as some positions. This is likely due to that the MSA was constructed using homologous proteins which fold to different structures, albeit similar. The design algorithm may have also introduced some bias.

**Fig 4.**
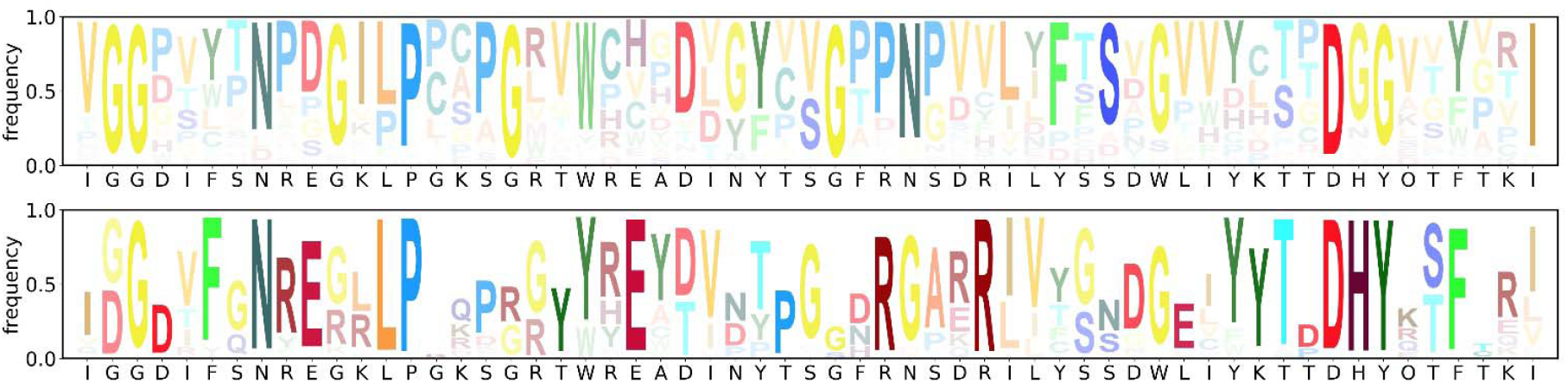
The logo plots for foldable designed sequences and multiple sequence alignment (MSA) of chain A of protein 1a2p. The part of the sequence with good alignment is shown (residues 49-107 out of total 108 residues). The figure with all the residues is provided in Supplementary material. The MSA of 1a2p has much stronger conservations than the foldable sequences. The average entropy for MSA and foldable designed sequences are 0.813 and 1.367, respectively. The numbers of residues with entropy smaller than 1 for MSA and foldable sequences are 65 and 29, respectively. Top panel: The logo plot for the foldable sequences whose predicted structures are within 3Å to the target structure. Bottom panel: The logo plot from multiple sequence alignment of 1a2p.

## Conclusion and Discussion

In this study, we used engineered and designed sequences to investigate the applicability of AlphaFold to problems other than structure prediction for naturally occurring sequences. We used mainly two types of sequences: point mutations with experimentally measured stability changes^18^, and sequences designed to fold to target protein structures using a modified algorithm based on ProDCoNN^11^. We found that the representations learned by AlphaFold during the prediction process can accurately predict the stability changes of point mutations. We also found that AlphaFold predicted the ProDCoNN designed sequences with a wide range of RMSDs to the target structures, indicating that AlphaFold can distinguish these designed sequences and some of them are more foldable than others. It also validated the effectiveness of ProDCoNN for the IPF problem. Using a modified ProDCoNN, we designed a framework for estimating the designability of protein structures, and for the first time estimated the designability of some natural proteins. Finally, comparing the foldable sequences for a target protein with its homologous sequences from a multiple sequence alignment showed significant differences between the two profiles. Studying such differences may shed light on the role of the conserved residues in the two profiles. Computational mutagenesis using AlphaFold starting from a natural sequence may help identify the minimum folding elements of the protein. Our findings in this study showed that several fundamental questions in computational structural biology can be immediately addressed with AlphaFold alone or combined with other previously developed methods.

### Protein engineering

Although the stability changes of point mutations cannot be directly inferred from the confidence scores of AlphaFold predictions, we found that the representations AlphaFold learned during the prediction process can be used to predict the stability change accurately. Several improvements may significantly increase the prediction accuracy. Firstly, the dataset we used can be substantially increased by using the data in a recent study^16^, which used more than 5000 point mutations. With more than doubled training data, we expect the model to have substantially improved performance; second, the “pair representation” generated during AlphaFold prediction should also be very useful for predicting stability changes. A challenge for using pair representation is that the dimensionality of the input will increase significantly, which may need to be regularized; third, in this study, we only took the information of the mutated residue from the single representation. Information of other residues will be very helpful to further improve the prediction performance. For example, we can take a fix number of residues from the sequence neighbors or spatial neighbors of the mutated residue. These improvements are being investigated by us currently.

### Estimating the designability of protein structures

Natural proteins have the capability to withstand a wide range of environmental stress and mutations. As seen from the point mutation data, a large number of mutated sequences of a protein can also fold to its native structure. The larger number of mutations a protein can tolerate, the more robust the protein structure and function is. The “designability” of a protein structure, defined as the number of sequences that encode that structure, has been proposed as an important property that contributes to the functional robustness of proteins^12^. Since protein structures can be organized as a hierarchy^17,19-21^ with four levels from folds to super families, to families, and to sequences, the designability of a protein fold is similarly defined as the number of families that take the fold as their native structures. A study has found that many disease-related proteins have folds with relatively few families, and a number of these proteins are associated with diseases occurring at high frequency^13^. This indicates that there is indeed a correlation between designability and functional robustness of proteins.

However, simply looking at the number of families under each fold is not a reliable measure for the fold’s designability and its functional robustness, because it has been found that the age of a fold correlates with its “usage” among natural proteins. For instance, eukaryotic folds found only in human, mouse, and yeast contain approximately 2.5 families, on average, compared to an average of 13.8 families per fold for all human proteins^13^. This observation has two implications: firstly, since the number of sequences exists in nature that can fold to a particular protein structure is not necessary a good indicator of its designability, we need to estimate the designability of a protein structure to have a better understanding of its functional robustness; second, since new folds have been much less “explored” by nature, there must exist new families, not related to any families found previously, that take one of the newer folds as their native structures. These new protein families may be hosts for some interesting, new functions. It is now possible to design new sequences using a protein design program such as ProDCoNN to specifically target on uncharted regions in the foldable sequence spaces and test the design with AlphaFold.

Since AlphaFold predicted that a significant portion of our designed sequences can fold to structures very close to the target structure, we formulated a framework for estimating the designability of a protein structure by combining a protein design algorithm, sequential Monte Carlo (SMC), and AlphaFold. To estimate designability, we need to sample foldable sequences using SMC. SMC is a special type of Monte Carlo method that allows one to estimate the partition function of a system^22^, which is usually very challenging to estimate. We have applied SMC in the past to estimate the entropy of lattice polymers^23^, the side chain entropy of proteins^24^, and other ensemble properties^25-27^. The total number of foldable sequences of a given protein structure can be expressed as a partition function, which can be estimated by SMC. In this study, for the first time, we have estimated the designability of a number of real proteins without unrealistic assumptions. As far as we know, designability has only been exhaustively enumerated for short chains using lattice models^12^ and small proteins (length 40 and 50) whose sequences were reduced to only two residue types (H and P)^28^.

### Predicting protein stability using natural and design sequences

As shown earlier, the representations learned by AlphaFold can be used to predict stability changes of point mutations. However, for a designed sequence which is much more different than any natural sequences, we still don’t know whether the structure predicted by AlphaFold is stable enough. Stability prediction for any given sequence is one of the key questions in protein folding unanswered by AlphaFold. Although predicting the exact stability can be quite challenging, it may be feasible to predict binary outcomes, such as whether a sequence has the stability similar to that of natural proteins. To address this with machine learning methods, we need to have stable sequences and unstable sequences. The protein sequences in PDB structures can serve as stable sequences. To obtain unstable sequences, one can randomly sample sequences, but these sequences are not challenging enough to train quality models to distinguish between foldable and unfoldable designed sequences. One reasonable option is to use all the designed sequences as negative sequences or use the designed sequences that are predicted to have RMSD to the target structure greater than certain threshold. With the sequence decoys and natural protein sequences, we can then train a model to perform a binary prediction: whether a sequence has stability comparable to natural proteins. When constructing the predictive models, we can again extract features from the representations learned by AlphaFold as the model input.

In protein design practice, we can select the sequences with small RMSDs to the target structure and optimize them by computational mutagenesis using a model for predicting stability changes for point mutations (e.g. the model we developed in this study). The stability of the optimized sequences can then be predicted by the binary model to check whether their stabilities are good enough. This provides a practical pipeline for designing sequences that can fold to the structures predicted by AlphaFold with satisfactory stability.

### Using AlphaFold to perform computational mutagenesis to understand the sequence-structure relationship of proteins

By studying the foldable sequences, we may identify residues that are important for the folding and stability of a protein structure and gain a deeper understanding of the sequence-structure relationship of proteins. This is a fundamental problem in structural biology. As defined earlier, the set of key residues for sequences to fold to a structure is called the minimum folding elements (MFEs) of the structure. Starting from foldable sequences or natural sequences, one can perform computational mutagenesis using AlphaFold to search for the MFEs for the protein structure. Each new sequence can be tested by AlphaFold for its foldability to keep updating the sets of foldable sequences. Stability prediction can also be performed to insure the sampled sequences are also stable. The search may eventually converge to certain sequence pattern which may serve as candidates for MFEs. Such studies may reveal some fundamental principles of protein folding and the sequence-structure relationship of proteins.

## Method and Data

AlphaFold and RoseTTAFold programs were obtained from their GitHub releases.

### Point mutations with experimentally measured stability changes

To study the correlation between AlphaFold confidence scores and the experimentally measured stability changes, we randomly selected 3507 experiments from protein single-point mutants stability database FireProtDB^18^, corresponding to 1251 mutants from 86 protein chains. The dataset contains 2557 experiments with Gibbs free energy changes (ΔΔ*G*) upon mutation and 952 experiments with changes in melting temperatures (Δ*Tm*). The stabilization status, defined by FireProtDB, fall into three categories: stabilizing mutations (Δ*Tm* > 1 or ΔΔ*G* < 1 kcal/mol), destabilizing (Δ*Tm* < 1 or ΔΔ*G* > 1 kcal/mol), and neutral (–1 ≤ Δ*Tm* ≤ 1 or – 1 ≤ ΔΔ*G* ≤ 1 kcal/mol). There are 328 stabilizing single-point mutants, 1842 destabilizing mutants, and 1337 neutral ones.

To train a model for predicting point mutation stability changes, we collected more data by randomly selecting 7777 experiments from FireProtDB with a valid ΔΔ*G*, corresponding to 2854 mutants from 114 protein chains. There are 499 stabilizing single-point mutants, 3653 destabilizing mutants, and 3625 neutral ones. We calculated the median of ΔΔ*G* if multiple experiments were performed for a particular point mutation, which results in 2854 data points, with 149 stabilizing mutants, 1311 destabilizing mutants, and 1394 neutral ones. The dataset is separated into 10 folds for 10-fold cross-validation. The separation is residue-based and guaranteed that two mutants at the equivalent site from two homologous proteins were always in the same fold. The homologous proteins are defined as sequence identity higher than 25%, which is calculated using T-coffee^29^.

### Designed sequences for inverse protein folding problem

We used a modified ProDCoNN^11^ to design sequences for 9 protein structures selected from SCOPe database^17^ which belong to 7 major classes defined by SCOPe. Specifically, two structures are from the *all alpha proteins* class (3lynA, 1e2aA), two structures from *all beta proteins* (1g6vK, 1kexA), one structure from each of the *alpha and beta (a/b)* (1h4yA), *alpha and beta(a+b)* (1a2pA), *multi-domain* (2avuF), *membrane and cell surface* (1g4yB), and *coiled coil proteins* (1ujwB) classes. The lengths of the proteins range from 76 to 156 residues. All the selected protein structures from PDB are single chains without missing residues and uncommon amino acids. No binding ligands were included.

The original ProDCoNN^11^ was not suitable for sampling protein sequences from the space of all possible sequences following certain probability distributions. To sample a sequence, we used a sequential Monte Carlo approach^22-26,30-32^ by sampling one residue at a time. A new residue is sampled conditioning on the residues sampled before it. This required us to train a sequential protein design model based on ProDCoNN. The sequential model starts from a backbone structure with no amino acid residues sampled for any positions and samples the amino acid type one residue at a time. While it samples an amino acid residue for a position, it conditions on the positions whose amino acid types have been sampled. The sampling order (which residue should be sampled the first, which residue the second, etc.) is decided based on the calculated entropy along the sequence:

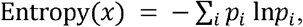

where *p*_*i*_ is the probability that residue *x* is predicted as amino acid type *i*. And the sampling probability is normalized by

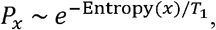

which includes a parameter temperature *T*_*1*_. The residues with lower entropy have a higher chance to be sampled first, while different sampling orders could be generated for different SMC samples.

When the sampling order is decided, the types of amino acids at each position will be sampled based on the predicted probabilities of the twenty amino acids by the trained ProDCoNN model. To adjust the relative probabilities of the types with non-zero probabilities, we used the following normalized probability:

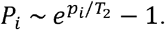

In our sampling, the parameter *T*_1_ was set to 1 and *T*_2_ was set to 0.05. The goals of tuning *T*_1_ and *T*_2_ are to have enough diversification in sampled sequences while also make sure a significant portion of the sequences are foldable. Because we used SMC to sample the sequences, it is expected that some sequences will not be foldable. The similarities between the sampled sequences and the wild type sequences range from 15.8% to 36%.

### Estimating the designability of a protein structure

To estimate the designability of a protein structure, we first used the SMC strategy described above to sample a number of sequences for the protein structure. We then used AlphaFold to predict the structures of these sequences. Those sequences with predicted structures with RMSD smaller than 4 Å to the target structure are considered as foldable and used for estimating the designability. The other sequences are discarded. Each sampled sequence using SMC has a weight, which is updated recursively as follows: *w*_*t*_ = *w*_*t*-1_/*p*_*t*_, where *w*_*t*_ is the weight at step *t, w*_*t-*1_ is the weight at step *t*-1 and *p*_*t*_ is the probability the actual amino acid type at step *t* is sampled. The designability can then be estimated using the equation: 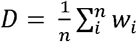, where *D* is designability, *w*_*i*_ is the weight of sequence *i*, and *n* is the total number of foldable sequences.

### Deep learning model for predicting stability changes of point mutations

We implemented a multilayer perceptron regression model for ΔΔ*G* and ΔΔ*Tm* prediction using features extracted from the representations learned by AlphaFold as input.

We first used AlphaFold to predict the structures of both wild type and mutant sequences. We then extracted the feature vectors from the position of the mutated residue from the “single representation” of the AlphaFold models for both wild type and mutant sequences as model input. We additionally include the subtraction of the two vectors from wild type and mutant as the input feature. As the dimension for each residue for the single representation is 384, the dimension of the final input feature vector is 3×384 = 1,152.

We built the model with four linear projections with input and output feature dimensions (1,152, 1,152), (1,152, 512), (512, 512), and (512, 1), and used the output in the last layer as the ΔΔ*G* (or ΔΔ*Tm*) value. Non-linear activation function ReLU was inserted between the linear projections. We used Adam optimizer and set the batch size as 1,024, learning rate as 1E-4, and training epochs as 1,000. The model was trained with Smooth L1 Loss.

### Definition of foldable sequences

In this study, we define a foldable sequence to a given structure as one that can fold to a structure with a RMSD smaller than 4Å to the given structure. Since we used AlphaFold to predict the structures of designed sequences, the foldability is predicted, not experimentally measured. In fact, the designed sequences, even predicted as foldable, may not have stabilities similar to real proteins. Our assumption here is that even a designed sequence is not actually foldable, there is a sequence in the neighborhood of it, which is actually foldable. Here the neighborhood of a sequence is defined as sequences with small number of mutations (i.e. smaller than 5) from the given sequence.

## Acknowledgement

JZ is supported partially by a grant from National Institute of General Medical Science of National Institutes of Health, grant # R01GM126558. Parts of the computational experiments were performed on a GPU cluster supported by an NSF infrastructure grant, OAC 1920147. The funder had no role in the study design, data collection and analysis, decision to publish, or preparation of the manuscript.

